# Proteomic characterization of aging-driven changes in the mouse brain by co-expression network analysis

**DOI:** 10.1101/2023.05.23.542020

**Authors:** Kazuya Tsumagari, Yoshiaki Sato, Hirofumi Aoyagi, Hideyuki Okano, Junro Kuromitsu

**Author notes:** **Correspondence:** (KT), (JK), /.

## Abstract

Brain aging causes a progressive decline in functional capacity and is a strong risk factor for dementias such as Alzheimer’s disease. To characterize age-related proteomic changes in the brain, we used quantitative proteomics to examine brain tissues, cortex and hippocampus, of mice at three age points (3, 15, and 24 months old), and quantified more than 7,000 proteins in total with high reproducibility. We found that many of the proteins upregulated with age were extracellular proteins, such as extracellular matrix proteins and secreted proteins, associated with glial cells. On the other hand, many of the significantly downregulated proteins were associated with synapses, particularly postsynaptic density, specifically in the cortex but not in the hippocampus. Our datasets will be helpful as resources for understanding the molecular basis of brain aging.

## Introduction

Aging is a time-dependent functional decline caused by the accumulation of cellular damage, that results in a progressive loss of physiological integrity, reduced function, and increased susceptibility to death (1). Brain aging causes a progressive decline in functional capabilities, resulting in impairments in learning and memory, attention, decision-making speed, sensory perception, and motor coordination (2). Examination of the brain at the cellular level has revealed various hallmarks of aging, including mitochondrial dysfunction, intracellular accumulation of oxidatively damaged proteins, dysregulated energy metabolism, loss of proteostasis, impaired adaptive stress response signaling, compromised DNA repair, aberrant neuronal network activity, dysregulated neuronal Ca^2+^ handling, and inflammation (1, 2). These age-associated changes in the brain are a strong risk factor for dementias such as Alzheimer’s disease (3), and thus a deeper understanding of the molecular basis of brain aging is a crucial task for elucidating the mechanisms of these diseases.

To date, several proteomics studies have addressed brain aging by analyzing model organisms. Walther and Mann analyzed mouse brain tissues using stable isotope labeling with amino acids in cell culture (SILAC). They accurately quantified more than 4,000 proteins, and demonstrated that proteome changes in brain aging are very small, at least at the bulk proteome level, and the proteome is robustly maintained to a relatively old age (4). Similar results have been obtained in studies by other groups. Ori et al. performed integrated transcriptome and proteome analyses and found that age-specific variations of the transcriptome and proteome are much less pronounced than tissue-specific differences (5). Yu et al. investigated nine mouse organs by targeted and non-targeted proteomics using isobaric tag quantitation and found that white adipose tissue is most affected by aging, while other tissues, especially brain, show very small proteomic changes (6). However, despite their small magnitude, changes in protein expression are likely to be closely associated with age-related cognitive decline (7), and importantly, these previous studies did not establish which function-related proteins change with aging in the brain. To this end, an approach is needed that can capture and interpret minute changes.

In this study, in order to generate large-scale proteomic datasets of mouse brain tissues with aging, we used a workflow involving tandem mass tag (TMT) quantitation, which is one of the most reproducible methods in quantitative proteomics, coupled with pre-fractionation and high-resolution mass spectrometry. We applied weighted gene co-expression network analysis (WGCNA) (8) to the dataset and detected co-expression modules associated with aging.

## Experimental procedures

### Mice

C57BL/6J Jcl male mice (CLEA Japan, Inc., Tokyo, Japan) were used in this study and maintained as described previously (9). Animal care and experimental procedures were performed in an animal facility accredited by the Health Science Center for Accreditation of Laboratory Animal Care and Use of the Japan Health Sciences Foundation. Experimental protocols were approved by the Institutional Animal Care and Use Committee of Eisai Co., Ltd., Tsukuba Research Laboratories.

### Experimental design and statistical rationale

Mice (*N* = 6 in each case) were sacrificed at 3, 15, or 24 months of age and perfused with phosphate-buffered saline, and then the cortices and hippocampi were collected. The hemispheres of the tissues were immediately frozen in liquid nitrogen and stored at −80°C. The right half of each tissue was utilized in this study. Statistical comparisons between two age groups were performed with Welch’s t-test employing a 5% permutation-based false discovery rate (FDR) filter.

### Sample preparation

Sample preparation was performed as described previously (9). Brain tissues were freeze-crushed using a multi-beads shocker (Yasui Kikai, Osaka, Japan). Proteins were extracted with a lysis buffer consisting of 4% SDS, 100 mM Tris-HCl (pH 8.5), 10 mM tris(2-carboxyethyl)phosphine, 40 mM 2-chloroacetamide, and HALT protease/phosphatase inhibitor cocktail (Thermo Fisher Scientific). Then, the proteins were purified by acetone precipitation and digested with LysC (FUJIFILM Wako, Osaka, Japan) and trypsin (Promega, Madison, WI). The resulting peptides were desalted on InertSep RP-C18 columns (GL Sciences, Tokyo, Japan) and TMT-labeled. The labeled peptides were fractionated by high-pH reversed-phase chromatography into 24 fractions.

### Nanoscale liquid chromatography/tandem mass spectrometry

NanoLC/MS/MS analysis was performed as described previously (9). The system consisted of an UltiMate 3000RSLCnano pump (Thermo Fisher Scientific) and an Orbitrap Fusion Lumos tribrid mass spectrometer (Thermo Fisher Scientific) equipped with a Dream spray electrospray ionization source (AMR Inc., Tokyo, Japan). Peptides were injected by an HTC-PAL autosampler (CTC Analytics, Zwingen, Switzerland), loaded on a 15 cm fused-silica emitter packed with 3 µm C18 beads (Nikkyo Technos), and separated by a linear gradient (5% solvent B for 1 min, 5−15% solvent B in 4 min, 15−40% solvent B in 100 min, 40−99% solvent B in 5 min, and 99% solvent B for 10 min; solvent A was 0.1% formic acid, and solvent B was 0.1% formic acid in 80% ACN) at the flow rate of 300 nL/min. All MS1 spectra were acquired over the range of 375–1500 m/z in the Orbitrap analyzer (resolution = 120,000, maximum injection time = 50 ms, automatic gain control = standard). For the subsequent MS/MS analysis, precursor ions were selected and isolated in top-speed mode (cycle time = 3 sec, isolation window = 0.7 m/z), activated by collision-induced dissociation (CID; normalized collision energy = 35), and detected in the ion trap analyzer (turbo mode, maximum injection time = auto, automatic gain control = standard). The top 10 most intense fragment ions were subjected to TMT-reporter ion quantification by SPS-MS3 (HCD normalized collision energy = 65) (10).

### Raw LC/MS/MS data processing

LC/MS/MS raw data were processed using MaxQuant (v.1.6.17.0) (11). Database search was implemented against the UniProt mouse reference proteome database (May 2019) including isoform sequences (62,656 entries). The following parameters were applied: precursor mass tolerance of 4.5 ppm, fragment ion mass tolerance of 20 ppm, and up to two missed cleavages. TMT-126 was set to the reference channel, and the match-between-run function was enabled (12). Cysteine carbamidomethylation was set as a fixed modification, while methionine oxidation and acetylation on the protein N-terminus were allowed as variable modifications. False discovery rates were estimated by searching against a reversed decoy database and filtered for <1% at the peptide-spectrum match and protein levels. Correction for isotope impurities was done based on the manufacturer’s product data sheet of TMT reagents.

### TMT-reporter intensity normalization

TMT-reporter intensity normalization among six of the 11-plexes was performed according to the internal reference scaling method (13) by scaling the intensity of the reference channel (TMT-126) to the respective protein intensities. Then, the intensities were quantile-normalized, and batch effects were corrected, using the limma package (v.3.42.2) in the R framework.

### Construction of weighted protein co-expression network

A weighted protein co-expression network analysis was performed using the WGCNA package (v.1.70-3) (8) in the R framework, as previously described (9). Expression similarities (adjacencies) were computed by the adjacency() function with the soft thresholding power of 26. The network type was set to “signed”. The adjacencies were transformed into a topological overlap matrix (TOM) by the TOMsimilarity() function, and a hierarchical clustering of proteins was performed by the flashClust() function (method = “average”) in the flashClust package (v.1.01-2), based on the corresponding TOM dissimilarity (1-TOM). Modules were detected by the cutreeDynamic() function using the hybrid tree cut method (deep split = 1, minimum module size = 50). Pearson correlations between each protein and each module eigenprotein, based on module memberships (kMEs) calculated by the moduleEigengenes() function, were calculated, and proteins without a significant correlation with the eigenproteins (p-value <0.05) based on the Pearson correlation were excluded from the module. Then, module eigenproteins were re-calculated and used for the downstream analyses.

### Statistics and bioinformatics analyses

Welch’s t-test and following permutation-based FDR calculation were performed using Perseus (14). Module-age correlations were calculated as Pearson correlations between the module eigenproteins and age, and q-values were calculated by the Benjamini-Hochberg method. GO term enrichment analysis and subsequent calculation of *q*-values by the Benjamini-Hochberg method were performed using R with the anRichment package. For cell-type-specific marker protein enrichment analysis, proteins that were at least 8-fold more highly expressed in a certain cell type than in other cell types in the dataset by Sharma et al. (15) were used as cell-type-specific marker proteins. Human UniProt accessions were converted to mouse accessions of the corresponding orthologs based on the HGNC comparison of orthology predictions (HCOP; https://www.genenames.org/tools/hcop/). Enrichment analyses for cell-type-specific marker proteins and cognitive stability-associated proteins were performed by means of the hypergeometric test, and q-values were calculated by the Benjamini-Hochberg method in the R framework. Protein-protein interaction analysis was performed using STRING (v.11.0) (16) (interaction sources = “Experiments” and “Databases”, minimum score = “medium confidence (0.400)”) and visualized using Cytoscape (v.3.8.0) (17).

## Results

### Deep and reproducible proteome profiling of the aging mouse brain tissues

We investigated proteomes of cortex and hippocampus dissected from mice at 3, 15, and 24 months old. In order to achieve large-scale and reproducible protein quantification, we employed a shotgun proteomics workflow consisting of TMT-labeling, fractionation by high-pH reversed-phase chromatography, and nanoLC/MS/MS (Fig. 1A). For each tissue, portions of each extracted protein were pooled, digested, labeled with TMT-126 or TMT-131C, and spiked as internal references for bridging two TMT-11-plexes and accounting for technical variation, respectively. In total, 6,821 and 6,910 proteins were quantified in at least three biological replicates in cortex and hippocampus, respectively, affording a total of 7,168 proteins (Fig. 1B). Reproducibility in protein quantification was good, with Pearson correlation coefficients >0.99 (Fig. 1C). Moreover, the median values of relative standard deviation (RSD) of protein quantification in the groups were less than 1% (Fig. 1D). Given the depth of the proteome and the inter-measurement or sample-to-sample reproducibility, we consider that our datasets are suitable for quantitative analysis of proteome alteration with aging.

**Figure 1.**
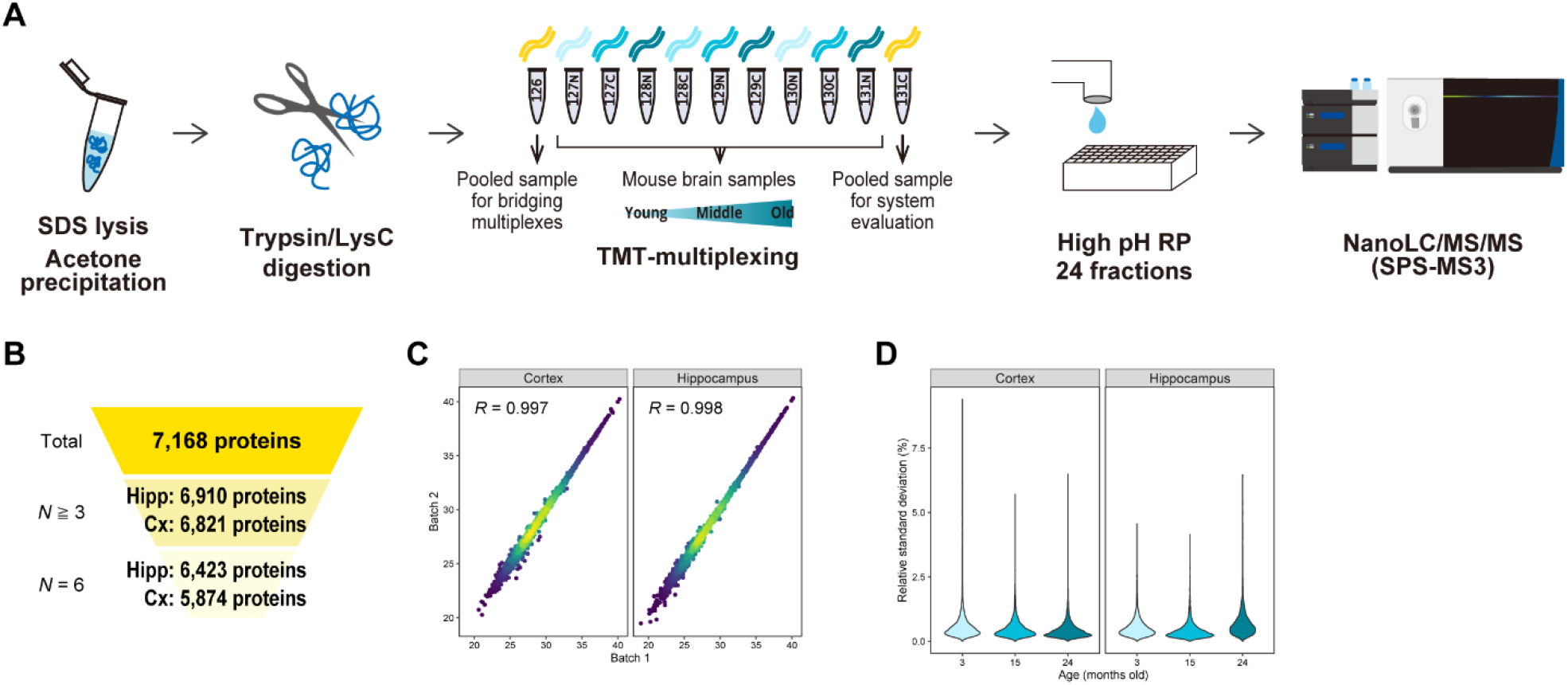
Deep and precise proteome profiling of mouse brain tissues with aging. **(A)** Proteins were extracted using sodium dodecyl sulfate (SDS), purified by acetone precipitation, and digested with LysC and trypsin. Six biological replicates were analyzed by six TMT-11-plexes for each of cortex and hippocampus. The digests were multiplexed by TMT, fractionated by high-pH reversed-phased chromatography (high pH RP), and analyzed by nanoLC/MS/MS. **(B)**Numbers of quantified proteins. *N*≧3, the number of proteins quantified in at least three replicates. *N*=6, the number of proteins quantified in all (six) replicates. Hipp, hippocampus. Cx, cortex. **(C)**Reproducibility of protein quantification between two measurements. Pearson correlation coefficients of the pooled samples (TMT-131C) are shown for cortex and hippocampus, respectively. **(D)**Reproducibility of biological replicates. Relative standard deviation (RSD) of each protein at the respective ages are shown for cortex and hippocampus.

### Extracellular proteins are upregulated, and synaptic function-related proteins are downregulated during aging specifically in cortex

We created volcano plots with a truncation at a FDR of 0.05 (Fig. 2A, B, S1). In the comparison between 3 and 24 months old, 133 and 52 proteins were significantly up- and downregulated in the cortex, while 150 and 93 proteins were significantly up- and downregulated in the hippocampus, respectively. These proteins accounted for only 2.7% and 3.5% of the total quantified proteins in the cortex and hippocampus, confirming that the change in the brain proteome driven by aging is very small (5, 6). In the two tissues, 47 proteins, including signal transducer and activator of transcription 1 (STAT1), myelin basic protein (MBP), glial fibrillary acidic protein (GFAP), and complement proteins C1qa, C1qb, and C4b, were commonly upregulated (Fig. 2C). Likewise, 11 proteins, including tenascin (TNC) and histone 1.5 (HIST1H1B), were commonly downregulated (Fig. 2D). Compared to the proteome change between 3 and 24 months old, the proteome changes between 3 and 15 months old and between 15 and 24 months old were smaller (Fig. S1). In the cortex, C4b and serine protease HTRA1 were found to be significantly upregulated both between 3 and 15 months old and between 15 and 24 months old, indicating that these proteins are progressively upregulated during aging. No such proteins were found in the hippocampus or in the upregulated proteins in the cortex. For the characterization of these significantly altered proteins, gene ontology (GO) term enrichment analysis was performed (Fig. 2E, F). The significantly upregulated proteins abundantly included extracellular proteins such as extracellular matrix proteins and secreted proteins. Among the downregulated proteins, significantly enriched terms were obtained only in the cortex, and these were related to synaptic functions.

**Figure 2.**
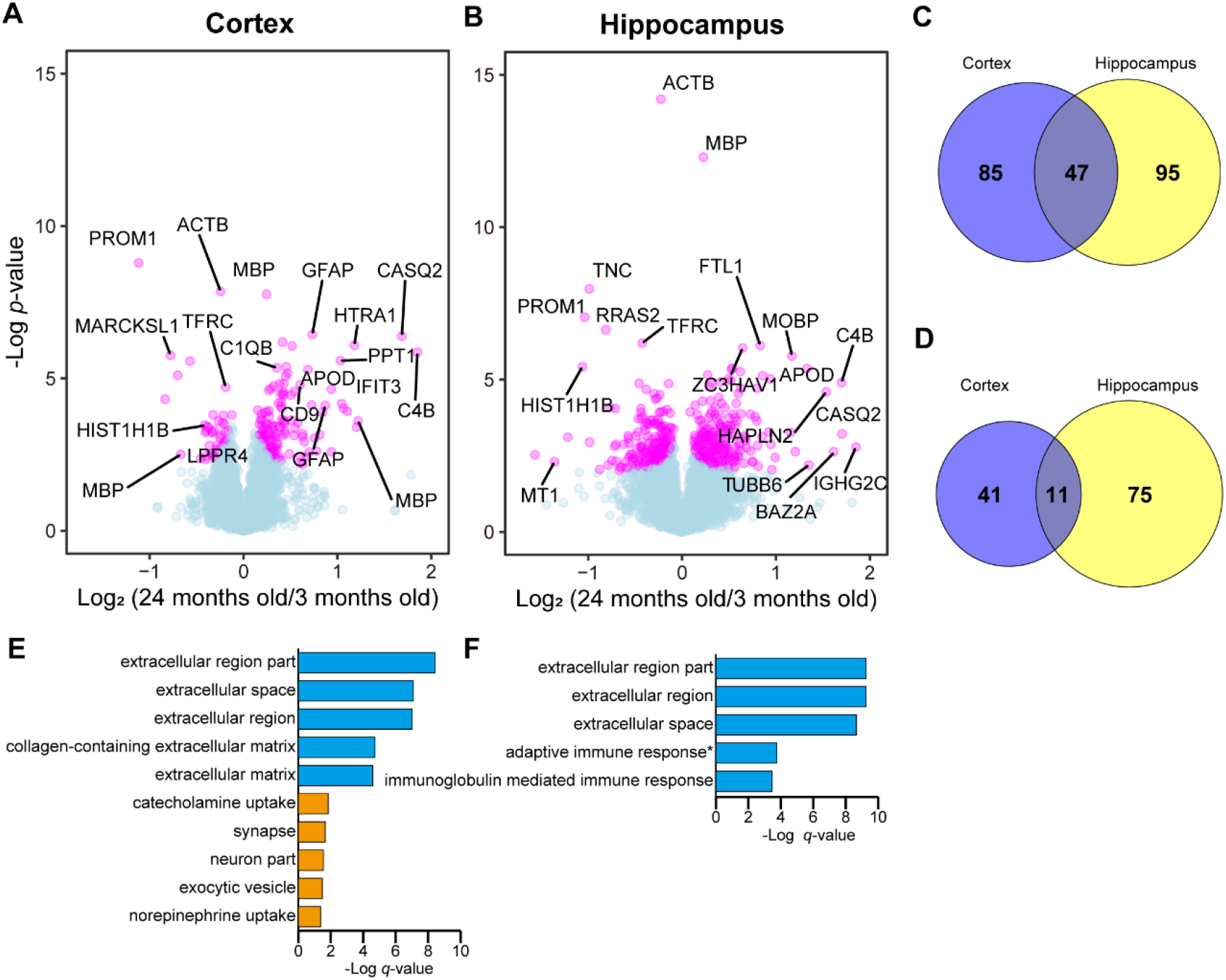
Comparison of protein expression at 3 months old and 24 months old. **(A, B)** Volcano plots comparing protein expression at 3 months old and 24 months old in cortex and hippocampus, respectively. Welch’s t-tests were performed to identify significantly changed proteins (N = 6). The proteins with q-value <0.05 are highlighted with color. Volcano plots comparing 3 months old and 15 months old, and 15 months old and 24 months old are shown in Figure S1. **(C, D)** Overlaps of significantly upregulated (C) and downregulated proteins (D) between cortex and hippocampus, among the commonly identified proteins in these tissues. **(E, F)** GO term (“molecular function” and “biological process”) enrichment analysis for significantly upregulated (blue) and downregulated proteins (orange) in cortex (E) and hippocampus (F). The top 5 terms are shown. No terms were significantly enriched for the downregulated proteins in hippocampus.

### Co-expression network analysis reveals age-associated protein modules

Co-expression network analyses, namely WGCNA, is a powerful and sensitive approach for the proteomic analysis of complex samples, such as brain tissues derived from patients with Alzheimer’s disease (18, 19) or disease models (9, 20). Such a sensitive analytical method is expected to be effective also for the interpretation of the changes in the brain proteome with aging. We applied the WGCNA algorithm to the cortex dataset and detected nine modules spanning diverse biological processes and cellular components based on GO terms (Fig. 3A, B). To find modules related to aging, we calculated the Pearson correlation coefficients between age and each module eigenprotein (Fig. S2), which is defined as the first principal component of a given module and serves as a representative (Fig. 3C, S2). The M1 synaptic module and M6 extracellular region module showed significant negative and positive correlations with aging, respectively, which again confirmed that synaptic proteins are downregulated, and extracellular proteins are upregulated during aging in the cortex. Notably, these modules abundantly included the significantly regulated proteins defined in the volcano plots. In the M1 synaptic module particularly, postsynaptic density proteins, such as HOMER1, DLGAP2 and 3, GRIN1 and 2B, and GRIA2, were abundantly included (Fig. S3). Moreover, consistently with this result, an enrichment analysis for cell-type-specific marker proteins (Fig. 3D) revealed that the M1 synaptic module was specifically enriched with neuronal marker proteins. In contrast, the M6 extracellular region module was enriched with glial proteins. Indeed, the M6 extracellular region module included many cell-marker proteins, such as GFAP (astrocytes) and MBP (oligodendrocytes). Consistent with the result of GO term enrichment analysis for the significantly upregulated proteins in the cortex (Fig. 2E), the M6 extracellular module also included ECM proteins such as collagen (type VI α-1, 3, and type XII α-1) and laminin (subunits α-1, 2, 5, β-2, and γ-1) proteins and secreted proteins such as complement proteins. Wingo et al. previously analyzed the dorsolateral prefrontal cortex (DPLFC) of human cohorts to investigate proteins that highly correlate with the cognitive trajectory and nominated 350 proteins that had increased abundance in cognitive stability (proteins with higher abundance in cognitive stability) and 229 proteins that had decreased abundance in cognitive stability (proteins with lower abundance in cognitive stability) (7). We asked how these proteins are regulated in the mouse cortex with aging (Fig. 3E). Interestingly, the proteins with lower abundance in cognitive stability showed highly significant enrichment in the M6 extracellular region module. For instance, GFAP, C4b, MBP, tight junction protein ZO-2, and N-Myc downstream regulated 1 (NDRG1) were included. The proteins with higher abundance in cognitive stability did not show significant enrichment for any of the modules. To address whether the detected co-expression networks were preserved in the hippocampus, we utilized module preservation statistics (Fig. 3F). Six modules detected in the cortex were significantly preserved with a Zsummary score >2. On particular, the M4 and M9 modules were highly preserved with a Zsummary score >10. On the other hand, three modules, including the M1 module, which was negatively correlated with aging, were not significantly preserved.

**Figure 3.**
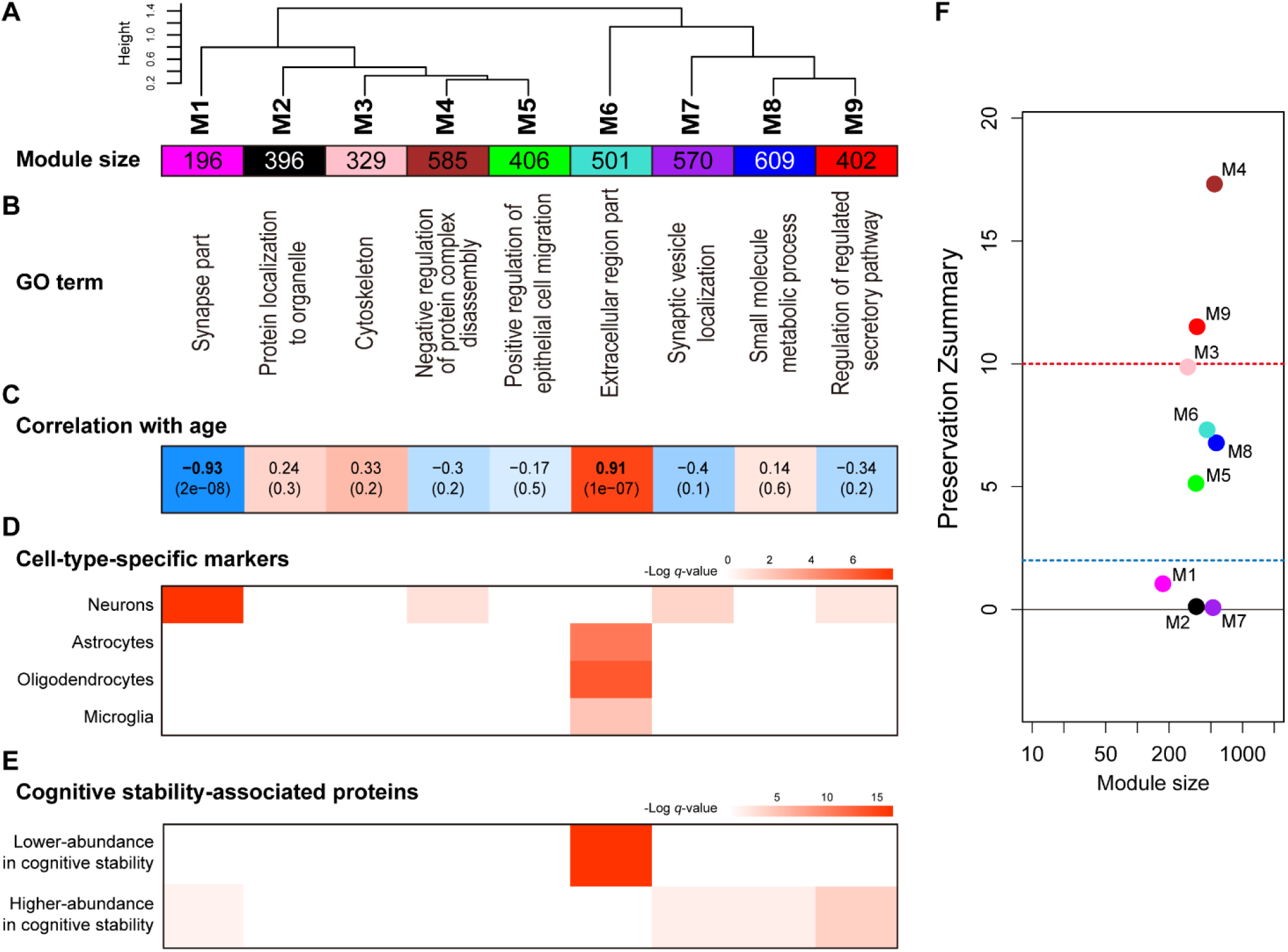
Weighted protein co-expression network analysis in aging cortex. **(A)**Module clustering dendrogram. Clustering was performed based on the eigenprotein values. **(B)**The most significantly enriched GO terms (“cellular component” or “biological process”). **(C)**Pearson correlation coefficients between each module eigenprotein value (Fig. S2) and age. The q-values are shown in parentheses. **(D)**Enrichment analysis for cell-type-specific marker proteins by hypergeometric test. **(E)**Enrichment analysis for cognitive stability-associated proteins by hypergeometric test. **(F)**Module preservation analysis examining whether the protein co-expression networks detected in the cortex proteomes were preserved in the hippocampus proteome. The dashed blue line indicates a Zsummary score of 2, above which module preservation was considered significant, and the dashed red line indicates a Zsummary score of 10, above which module preservation was considered highly significant.

## Discussion

Here, we generated protein expression datasets of the cortex and hippocampus in mice across three age points using a workflow consisting of sample multiplexing by TMT, pre-fractionation by high-pH reversed-phase chromatography, and high-resolution mass spectrometry. The quality of the data sets was validated by the high reproducibility of the measurements and the low RSD values. First, we identified significantly altered proteins using volcano plots. Then, we constructed a co-expression network based on the WGCNA algorithm and discovered two age-related modules. These two analyses consistently showed that many of the significantly upregulated proteins were extracellular proteins, while the significantly downregulated proteins, which were specifically observed in the cortex, were associated with synaptic functions.

The upregulated extracellular proteins, including ECM proteins such as collagens and laminins, as well as secreted proteins such as complement proteins, were associated with glial cells, as indicated by WGCNA. One of the important roles of laminin proteins in the brain is the organization of the blood-brain barrier (BBB) (21). Astrocyte-derived laminin is particularly important for the maintenance of BBB integrity (22). It is known that prolonged vascular flow on the basement membrane eventually leads to basement membrane thickening (23). Given this fact, the upregulation of the laminin proteins may reflect the thickened basement membrane at the neurovascular units. In addition, collagen type IV accumulates in the basal lamina of human cerebral microvessels with age (24). Taken together, the changes in the ECM proteins with aging may be associated with the alteration of the neurovascular system. Upregulation of complement proteins is tightly associated with neuroinflammatory events and has been observed in various neurodegenerative diseases, including Alzheimer’s disease and related disease models (9, 25, 26). Notably, GFAP, a marker protein of activated astrocytes (27), was significantly upregulated with aging and was incorporated into the M6 extracellular region module. We also found that myelin sheath proteins, including MBP, were significantly enriched in the M6 module. Oligodendrocytes remain active during aging, and it has been established that there is a substantial increase in the numbers of oligodendrocytes over the life span of monkeys (28). In summary, much of the protein upregulation in the brain is a composite result of the changes in different glial cells. In particular, the M6 extracellular region module was associated with cognitive decline, and so these proteins should be of interest in future studies.

We found that synaptic proteins, specifically postsynaptic density proteins, were downregulated by aging in the cortex. Most of the postsynaptic density proteins in the M1 synaptic module were not identified as significantly downregulated proteins in the volcano plot analysis, highlighting the sensitivity of our approach. Intriguingly, the M1 synaptic module was not significantly conserved in the hippocampal proteome. Proteomics changes in the levels of postsynaptic density proteins were also observed in a tauopathy mouse model (9). Assuming that the hippocampus is relatively resistant to the downregulation of synaptic proteins, elucidation of the underlying mechanisms could lead to the development of treatments for neurodegenerative diseases.

In conclusion, our deep and precise proteomic analysis allowed us to characterize age-related changes in the brain proteome. We believe that our dataset, together with the co-expression network, provide a basis for further studies to unravel the underlying mechanisms of brain aging, as well as age-related diseases such as Alzheimer’s disease.

## Acknowledgements

We would like to thank members of Eisai-Keio Innovation Laboratory for Dementia for fruitful discussions. KT is grateful to Koshi Imami (RIKEN, Yokohama) for supporting preparation of the manuscript at the current laboratory.

## Data Availability

The raw data and analysis files have been deposited to the ProteomeXchange Consortium via the jPOST partner repository (29) with the data set identifier PXD PXD041485 (JPST JPST001514).

## Supplemental Data

This article contains supplemental data. Supplementary Figures (.pdf)

Figure S1. Volcano plots comparing protein expression at 3 months old and 15 months old, and at 15 months old and 24 months old.

Figure S2. Levels of module eigenproteins.

Figure S3. Interactome of M1 synaptic module proteins.

## Author contributions

K.T. performed all experiment and downstream analyses and wrote the manuscript. Y.S, H.A, H.O. and J.K. supervised the study and reviewed the manuscript.

