## Supporting information for "Proteomic characterization of aging-driven changes in the mouse brain by co-expression network analysis"

### Contents

Supplementary Figures (.pdf)

- Figure S1. Volcano plots comparing protein expression at 3 months old and 15 months old, and at 15 months old and 24 months old.
- Figure S2. Levels of module eigenproteins.
- Figure S3. Interactome of M1 synaptic module proteins.

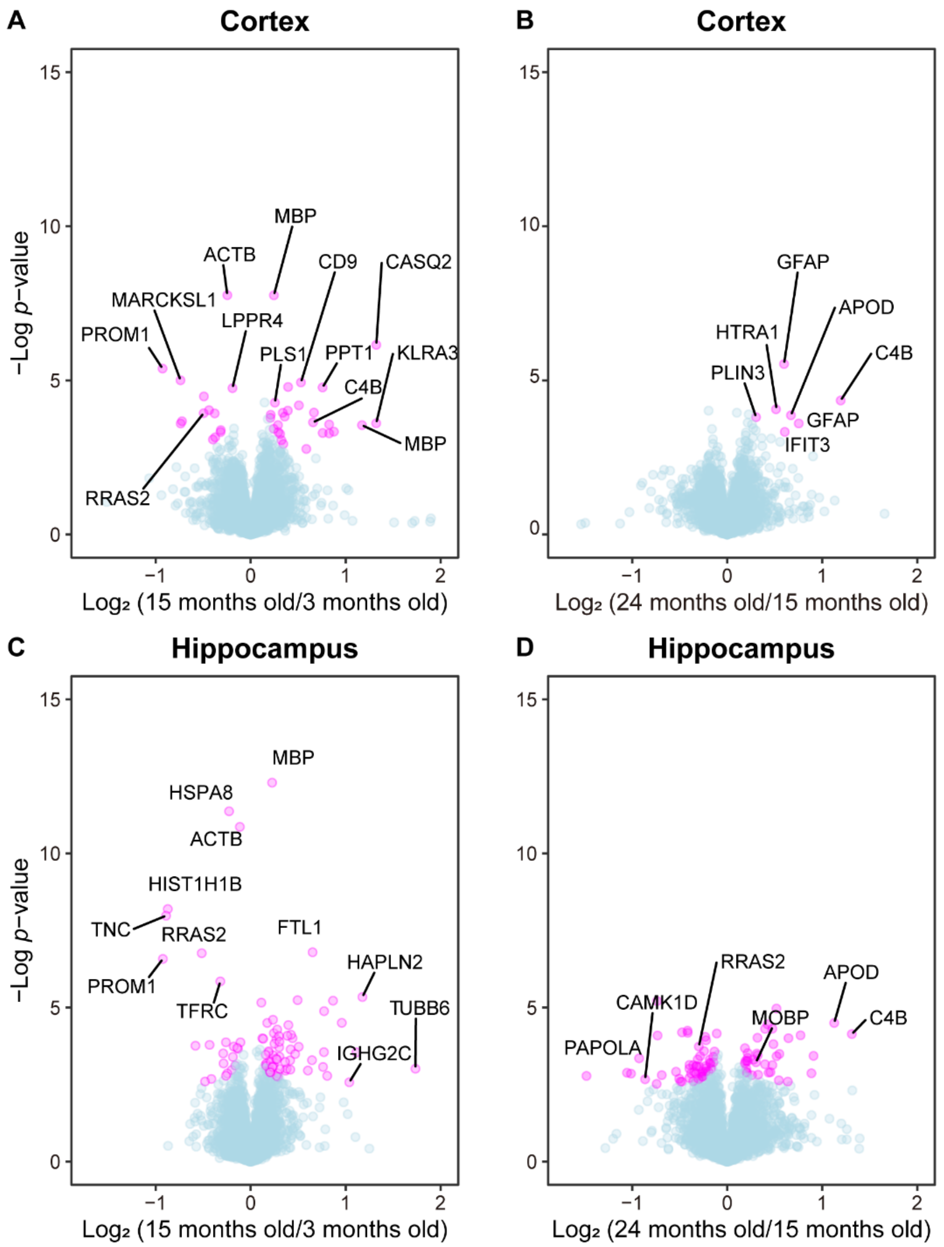


**Figure S1. Volcano plots comparing protein expression at 3 months old and 15 months old, and at 15 months old and 24 months old.**

Volcano plots comparing protein expression at 3 months old and 15 months old (A and C), and at 15 months old and 24 months old (B and D) in cortex (top row) and hippocampus (bottom row). Welch's t-tests were performed to identify significantly changed proteins ($N$ = 6). The proteins with $q$-value <0.05 are highlighted with color.

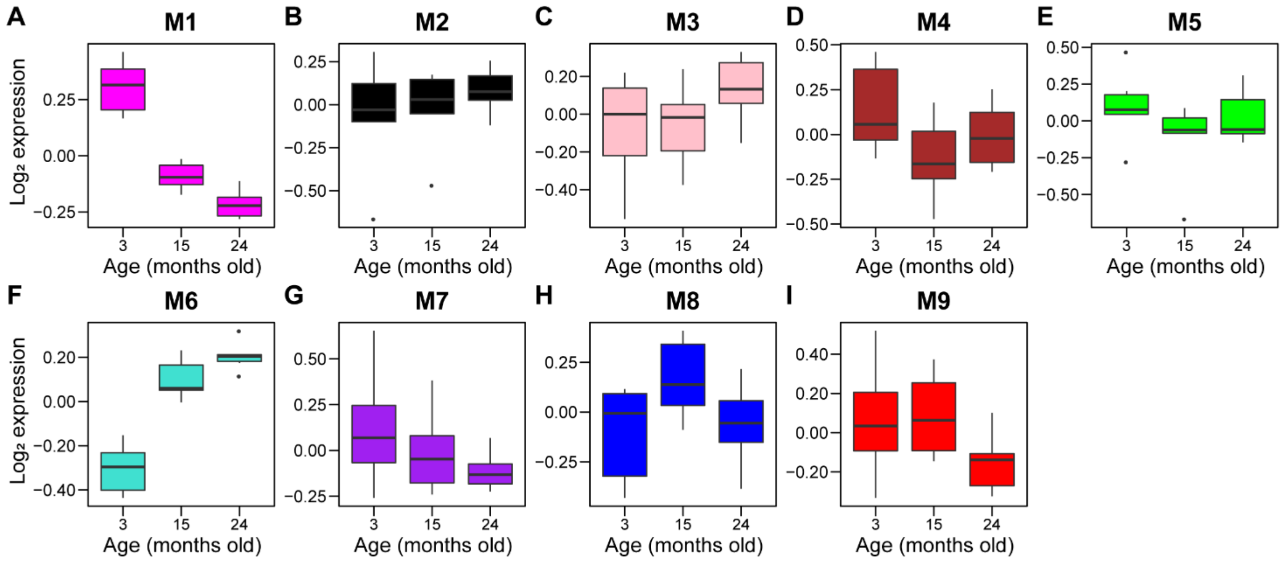


**Figure S2. Levels of module eigenproteins.**

Module eigenprotein is defined as the first principal component of a given module and serves as a representative of the module.

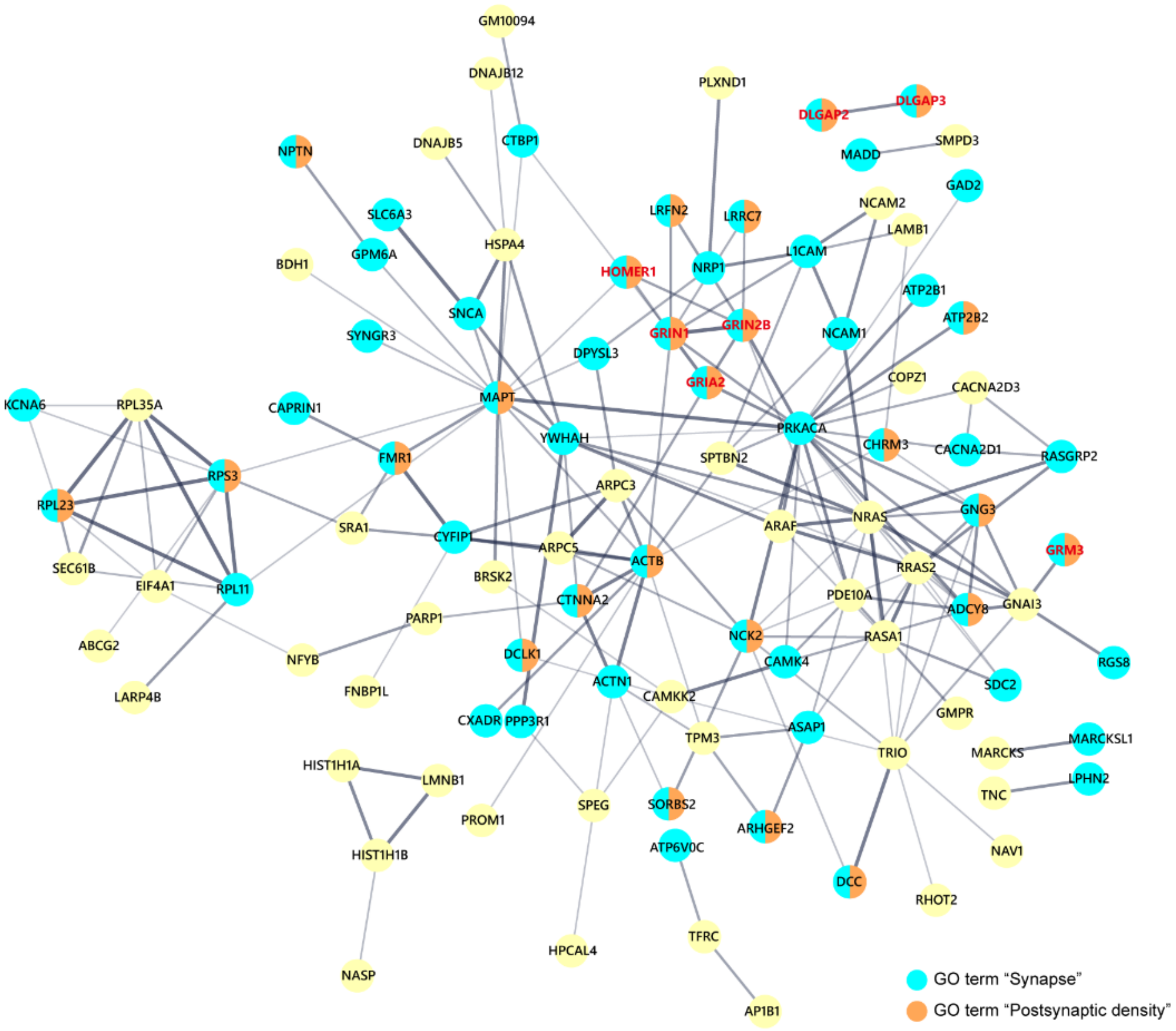


**Figure S3. Interactome of M1 synaptic module proteins.**

Protein-protein interaction of M1 synaptic module proteins. The proteins with GO terms “synapse” and “postsynaptic density” are highlighted in blue and brown, respectively.
